# Engineering self-organized criticality in living cells

**DOI:** 10.1101/2020.11.16.385385

**Authors:** Blai Vidiella, Antoni Guillamon, Josep Sardanyés, Victor Maull, Nuria Conde-Pueyo, Ricard Solé

## Abstract

Complex dynamical fluctuations, from molecular noise within cells, collective intelligence, brain dynamics or computer traffic have been shown to display noisy behaviour consistent with a critical state between order and disorder. Living close to the critical point can have a number of adaptive advantages and it has been conjectured that evolution could select (and even tend to) these critical states. One way of approaching such state is by means of so called self-organized criticality (SOC) where the system poises itself close to the critical point. Is this the case of living cells? It is difficult to test this idea given the enormous dimensionality associated with gene and metabolic webs. In this paper we present an alternative approach: to engineer synthetic gene networks displaying SOC behaviour. This is achieved by exploiting the presence of a saturation (congestion) phenomenon of the ClpXP protein degradation machinery in *E. coli* cells. Using a feedback design that detects and then reduces ClpXP congestion, a *critical motif* is built from a two-gene network system, where SOC can be successfully implemented. Both deterministic and stochastic models are used, consistently supporting the presence of criticality in intracellular traffic. The potential implications for both cellular dynamics and designed intracellular noise are discussed.

## I. INTRODUCTION

In order to adapt to environmental challenges, biological systems exhibit a diverse array of response mechanisms grounded in sensors and actuators as well as information-processing units. Adaptive responses require dynamical features that combine low energetic costs along with fast changes to efficiently respond to environmental changes. Flocks of birds and fish schools widely fluctuate in time but rapidly reorganize when a perturbation (such as the presence of a predator) occurs. Within cells, noise was early identified as playing multiple roles affecting cell fate, population heterogeneity, signal amplification or response to stress [1–3]. Noise is both an inevitable outcome of stochastic molecular interactions and an essential ingredient in decision making [4].

It has been shown that many complex systems seem to be poised close to so called critical points separating ordered from disordered states [5–8]. In a nutshell, both living and non-living systems organize at the boundary separating regular (predictable) from random (disordered) behaviours. At this point, complex dynamics with scale-invariant properties emerge [9, 10]. If *s* defines the total activity in one given event, such as number of firing neurons [11–14], gene expression [15–18], number of active ants in a colony [19, 20], critical epidemic bursts [21] or the size of traffic jams [22–24], the resulting distribution *P* (*s*) is a fat-tailed one, following a power-law of the form *P* (*s*) ∼*s*^−*γ*^, with a scaling exponent *γ* usually located within the interval 2 ≤ *γ* ≤ 3 [8, 25].

Critical points can be reached by fine-tuning a given “control” (or bifurcation) parameter *η* (Fig. 1a-b). This parameter (such as density of particles, temperature or reaction rate) directly influences the system’s state, as described by the *order parameter S* (system’s activity, for example). The way to criticality based on tuning key parameters is well illustrated by enzymatic queueing processes [26]. These authors used the framework of queueing theory (QTH) to study the dynamics of different proteins (the ‘customers’ in QTH) that are processed by a downstream set of enzymes that play the role of ‘servers’. Specifically, they considered the native *E. coli* protease, ClpXP. The key concept here is that the protease complexes are a limited resource (hereafter we will thus consider the amount of ClpXP as a constant) that can only ‘process’ (degrade) a limited number of incoming proteins. In Fig. 1a we provide a basic diagram considering a protein *σ* being expressed at some given rate *η*. If the rate of protein production is low (queues are short), degradation is efficient since the proteases can process all incoming *σ* units (free phase). If production is too high, a long queue of molecules ‘waiting’ to be processed will be present (congested phase). The two regimes are separated by a critical point where an optimal balance is reached, along with wide fluctuations in concentrations [26].

**FIG. 1:**
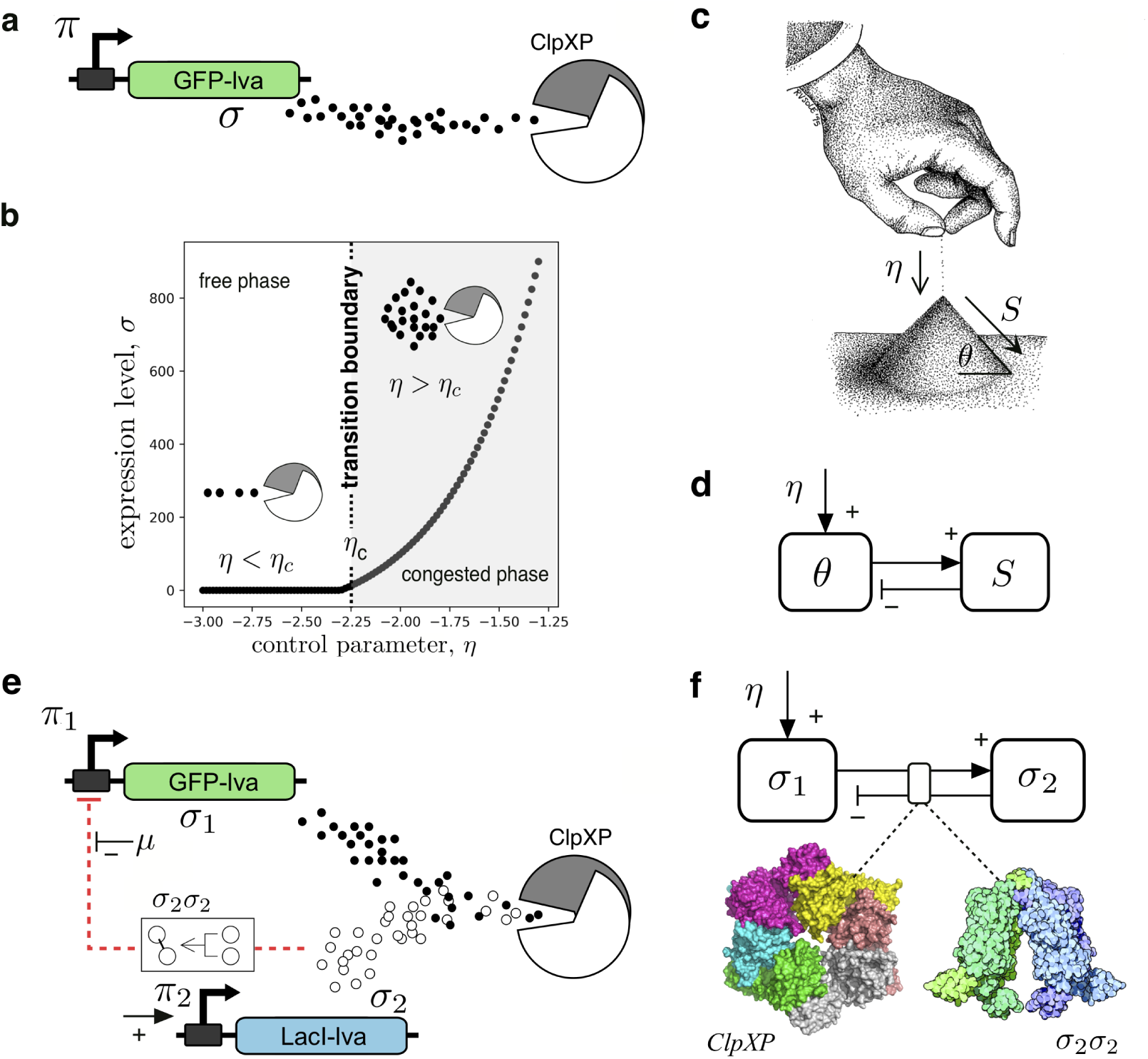
Paths to intracellular criticality. Tunable critical dynamics can be found in simple genetic circuits (a) where a given gene is constitutively expressed into a protein *σ* that decays and is also actively degraded by cell proteolytic machinery (ClpXP). By tuning expression rate *η* (d), a critical rate *η*_*c*_ is found to separate a phase of efficient degradation from another (light gray) involving congestion. In (b) the thick line indicates that few proteins are found for *η < η*_*c*_ (the proteolytic machinery efficiently degrades it) while it accumulates on the right side, due to congestion (ClpXP fails to degrade all the incoming proteins). An alternative, non-tuned path is self-organised criticality (SOC) which can emerge is provided by the sandpile (c). As grains of sand are slowly added at a rate *η*, the angle of the pile *θ* grows and only small avalanches will be observed. However, as the critical (maximum) *θ*_*c*_ is reached, avalanches of all sizes take place, reducing *θ*. The feedback between the *order parameter S* and the *control parameter θ* is summarised in (d). To facilitate the conditions enabling SOC, a two-gene circuit with negative feedback (e) allows mapping the sandpile feedback diagram (f). Here, both proteins compete for ClpXP (higher levels of *σ*_1_ also implies high values of *σ*_2_) and a repression feedback is mediated by *σ*_2_*σ*_2_ (the Lac repressor dimer) with *σ*_1_ and *σ*_2_ acting as order and control parameters, respectively.

The critical point is a rather unique one. Can these systems poise themselves into critical states with-out fine tuning? An alternative mechanism to reach critical states is provided by self-organized criticality (SOC) [27–29]. In this case, control and order parameters interact in such a way that the system spontaneously self-organizes into a critical state [30, 31]. The canonical example of SOC is the critical sandpile (Fig. 1c). By slowly adding grains of sand to the pile (at a rate *η*), its slope *θ* increases. At the beginning, only a few grains will fall down but the number *s* of grains in an avalanche rapidly grows as the angle of repose *θ*_*c*_ is approached. Once we have *θ* = *θ*_*c*_, the interaction between *θ* and sand avalanches (the order parameter) will keep the system at criticality [29]. This is summarized in Fig. 1d where the nature of the feedback between control and order parameters is sketched. Is a SOC mechanism a possible way of generating critical gene expression dynamics? In this paper we propose a novel approach to create SOC dynamics by engineering the interaction between order and control parameters in a simple two-gene network design.

## II. RESULTS

### A. Two-gene SOC motif model: deterministic and stochastic dynamics

The importance of the queueing dynamics in the enzymatic processing is illustrated by the *E. coli* stress response to starvation, which is triggered by an excess of mistranslated proteins. Stress can cause a significant increase in the concentration of aberrant proteins, which must be degraded. When such an overload occurs, the concentration of the sigma factor (the master stress regulator) builds up, eventually triggering the stress response [34]. Recent theoretical work also suggests that queueing could be adaptive in parallel enzymatic networks when the input flux of substrates is balanced by the maximum processing capacity of the network [35]. Here we go a step further and show how a simple SOC circuit can be actually engineered *in vivo*.

Our goal in this work is to define the basic design principle to build a “genetic sandpile” system that can capture the feedback structure shown in Fig. 1e. First of all, consider the simple, two-gene network circuit shown in Fig. 1e. Two proteins *σ*_1_ and *σ*_2_ resulting from their expression will be used as the building blocks for the order and control parameters, respectively, thus implementing a SOC feedback loop (Fig. 1f). The aim is to exploit the topology of the gene-gene interaction in such a way that the system can detect the degree of congestion of the ClpXP system by using *σ*_2_ as a sensor of the *σ*_1_ levels. Our control protein *σ*_2_ can form dimers, i. e. *σ*_2_ + *σ*_2_ *→ σ*_2_*σ*_2_ and this dimeric forms act as inhibitors (see Methods, eqs. (1-3)). If congestion occurs, the abundance of *σ*_2_ increases and its negative feedback effects also do so. A standard Hill function will be used to model the dimers as transcriptional repressors. The construction of our circuit involves two steps: (a) engineering the “critical” motif and (b) adjusting the protein production levels. This might seem contradictory with the “self-organized” description of SOC, but an example of why this is required is given by rice piles [36] which exhibit SOC but for some given grain aspect ratios.

Along with the topology of the SOC motif, a separation of scales is known to be a characteristic of SOC dynamics [31]. While the control parameter has a slow dynamics (the angle of the sandpile) the system’s response (the avalanche time scale) is fast. In our model, two additional parameters are used to favor the presence of criticality. These are the promoter efficiency for *σ*_2_, labelled *η*_2_; and an extra inhibition acting on the repressor *σ*_2_*σ*_2_ indicated as *µ* in Fig. 1e. We need to remember that degradation (and other dissipative events) affects *σ*_2_ and thus a minimal concentration of this sensor is needed in order to effectively detect congested states. On the other hand, in order to experimentally validate our model, we need to tune the strength of the feedback (required to trigger a rapid decay of the intracellular concentration of *σ*_1_). By tuning these two parameters, we include both the non-SOC design based on queueing as a special case [26] (see Section I in the SM) and a mechanism to achieve the SOC state.

In Fig. 2 we summarise the behaviour of the two-gene system (Fig. 1e; see Methods and Section II in the SM for mathematical details) using both deterministic (Section II.A, SM) and stochastic (Section II.B, SM) dynamics. A unique stable equilibrium (*σ*_*eq*_ = (*σ*_1,*eq*_, *σ*_2,*eq*_), indicated with a solid black circle in the (*σ*_1_, *σ*_2_) phase portraits of Fig. 2) is found, with a characteristic structure of the orbits in the phase space, as shown in Fig. 2a for the unregulated domain (here *η*_2_ = 10^−3^). As the expression rate *η*_2_ of the control parameter increases close to *η*_2_ ∼10^−2^, it is easy to see the presence of a slow-fast dynamics in the distinct structure of the vector fields consistently with the SOC requirement of time scale separation. Here the critical motif allows for large fluctuations in *σ*_1_ to occur (Fig. 2b) as shown by the compression of the trajectories in the phase portrait close to the fixed point, to be compared with the more homogeneous flow displayed in Fig. 2a for *η* = 10^−3^. The analysis of this system shows that, once close to the equilibrium point, small changes in the control *σ*_2_ trigger marked population spikes in *σ*_1_ (see Fig. S4, SM). Larger values for *η*_2_ (Fig. 2c) do not exhibit such a time scales separation. The analytic and numerical investigation of the eigenvalues of the fixed point to study its stability properties reveal the presence of a maximum in the so called ratio index (Fig. S7, SM), indicating a remarkable change in the vector field of the phase portrait when *η*_2_ ∼10^−2^ (given our set of fixed parameters indicated in Fig. 2).

**FIG. 2:**
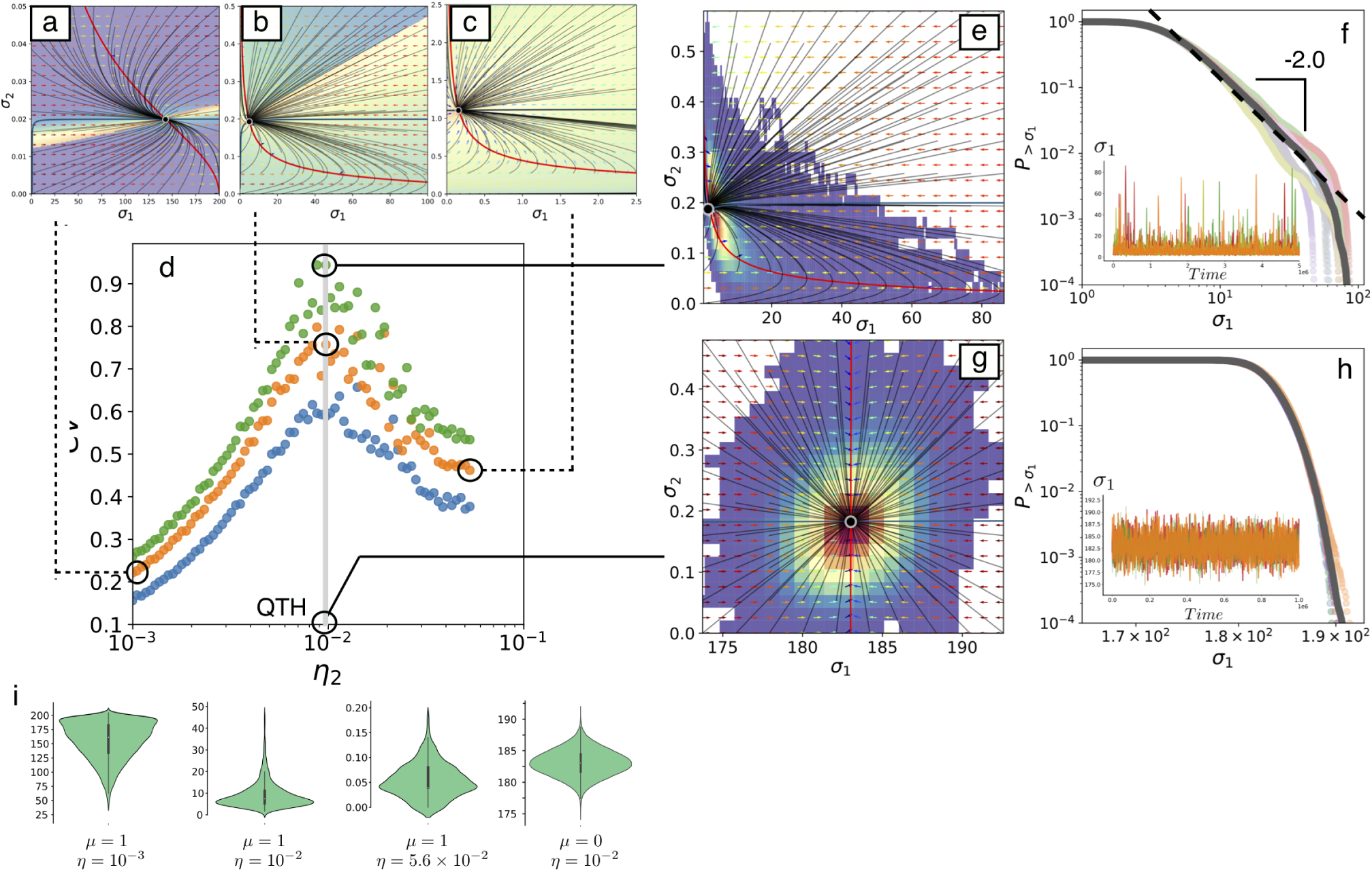
Nonlinear dynamics modeling of a two-gene critical motif. Following the causal scheme sketched in Fig.1 a-c, and defined by equations (1-2) in the Methods section, the trajectories of the system in the (*σ*_1_, *σ*_2_) space are displayed for increasing values of *η*_2_. Here the other parameters are *η*_1_ = 10^−2^, *δ* = 5 ×10^−2^, *δ*_*Clp*_ *C* = 10^−2^, *K* = *θ* = 10^−3^. The feedback control parameter is (a) *η*_2_ = 10^−3^, (b) *η*_2_ = 10^−2^ and (c) *η*_2_ = 0.056, respectively. The nullclines are plotted in red (*dσ*_1_*/dt* = 0) and blue (*dσ*_2_*/dt* = 0). The orbits (shown with black curves) flow towards a single attractor (*σ*_*eq*_). The vector field is indicated by unitary arrows, the color of which corresponds to their module (blue for small and red for the heavier). A background color scale is also used showing the arrival time required for each initial condition (*σ*_1_(0), *σ*_2_(0)). Here yellow and violet indicate short and long arrival times, respectively. In (b) a combination of slow decays in *σ*_2_ and fast responses for *σ*_1_ are at work. The stochastic dynamics of the model reveals a maximum in the *CV* when *η*_2_ ∼10^−2^, as shown in panel (d) where the colours stand for *µ* = 0.5 (blue), *µ* = 1.0 (orange) and *µ* = 1.5 (green). The relative location of the deterministic flows are indicated by dashed lines. Three different values of the coupling parameter *µ* are also used to show the robust nature of the maximum, where the SOC motif has been tuned to generate fat-tailed behaviour, as shown in panel (e). In this panel the hot map is plotted on top of the phase space, showing a maximum close to the attractor as well as the fat-tailed scaling behaviour (f) with 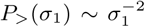 which gives *γ* ∼ 3 for *P* (*σ*_1_). Here five different runs are shown along with their average (dark line). By contrast, the flows and hot maps for the non-SOC circuit close to the queuing theory (QTH) transition (g) have a Gaussian pattern (h) with exponential tails as shown by the straight lines in the linear-log insets. The complete distribution is depicted with the upper violin plots (i) for different parameter values (as indicated). The effect on the coupling (*µ*) can be also observed in the different plotted simulation points.

To see how this nonlinear flows behave under the presence of intrinsic noise, a stochastic numerical implementation of the two-gene circuit has been carried out using the Gillespie method [32]. In Fig. 2d the coefficient of variation 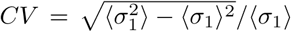 of the generated time series is displayed against *η*_2_ for three values of *µ*. This coefficient provides a statistical estimate of the variance of the fluctuations and a well-defined maximum is observed when *η*_2_ ∼10^−2^. In Fig. 2e we have overlapped several stochastic re-alisations with the vector field close to the maximum *CV* (for *η*_2_ = 10^−2^). The density plot reveals that the stochastic system visits very frequently the fixed point *σ*_*eq*_ (orange-red colours in Fig. 2e), but also wanders far away in the lower part of the phase portrait, where the vector field is faster and pushes the stochastic paths far away from the equilibrium point, then returning it back to the deterministic equilibrium. In Fig. 2f the resulting distribution is displayed. Specifically, if *P* (*σ*_1_) indicates the probability distribution of *σ*_1_ expression levels (activity), the cumulative dis-tribution is defined as 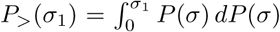 and helps smoothing the random noise exhibited by *P* (*σ*_1_). If the original distribution follows a scaling 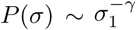, the cumulative one gives *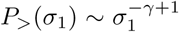*. The stochastic model gives a value of *γ* ∼ 3 (fig. 2f) es-timated from the average of five different runs. The time series associated to this parameter combination is shown in the inset, revealing a characteristic bursting dynamics typical of the SOC state. These results can be compared with the smooth and Gaussian behaviour obtained with the unregulated dynamics setting *µ* = 0 (Fig. 2g-h).

### B. Engineering a synthetic SOC circuit in *E. coli*

The theoretical model predicts that the SOC feedback loop defined above (Fig. 1e-f) will display bursting dynamics with fat-tailed activity distributions associated to the *σ*_1_ protein (order parameter). If the expression level (*η*_2_) of the control protein *σ*_2_ is large enough, its concentration will act as a congestion sen-sor and will repress *σ*_1_, following the SOC motif design. Otherwise, the system will lack the feedback loop. Here we show how can we engineer the genes circuit incorporating some parameters that allow including the non-SOC phase transition as described above including intracellular queueing processes as a special case. Since the time scale of expression changes is comparable to the replication rate of individual cells, no individual time series can be gathered, but instead the collective response of the system will be analyzed to detect the presence of a SOC state. The underlying assumption is thus that we have a colony-level sampling of the dynamical states and the critical dynamics is assessed by looking at the distribution of cell states and the resulting aggregated statistics (as-suming ergodicity).

The explicit experimental design of our SOC circuit implementation is outlined in Fig. 3a-b. The order parameter is encoded by the green fluorescent protein (GFP), and the control parameter acting as the congestion sensor, by the LacI repressor protein, the relative expression of which will be estimated by means of the expression of the red fluorescent protein (RFP). In both cases we use the unstable variants GFP-lva and RFP-lva, respectively. The construct expresses GFP under the pLacI promoter (*η*_1_) while the LacI repressor and RFP reporter protein are under the pBAD promoter, with non leaky tight regulation and highlevel expression inducible by Arabinose (*η*_2_) [52–54]. All three proteins of the circuit are tagged with lvasequence to be degradable by the ClpXP proteolytic complex.

**FIG. 3:**
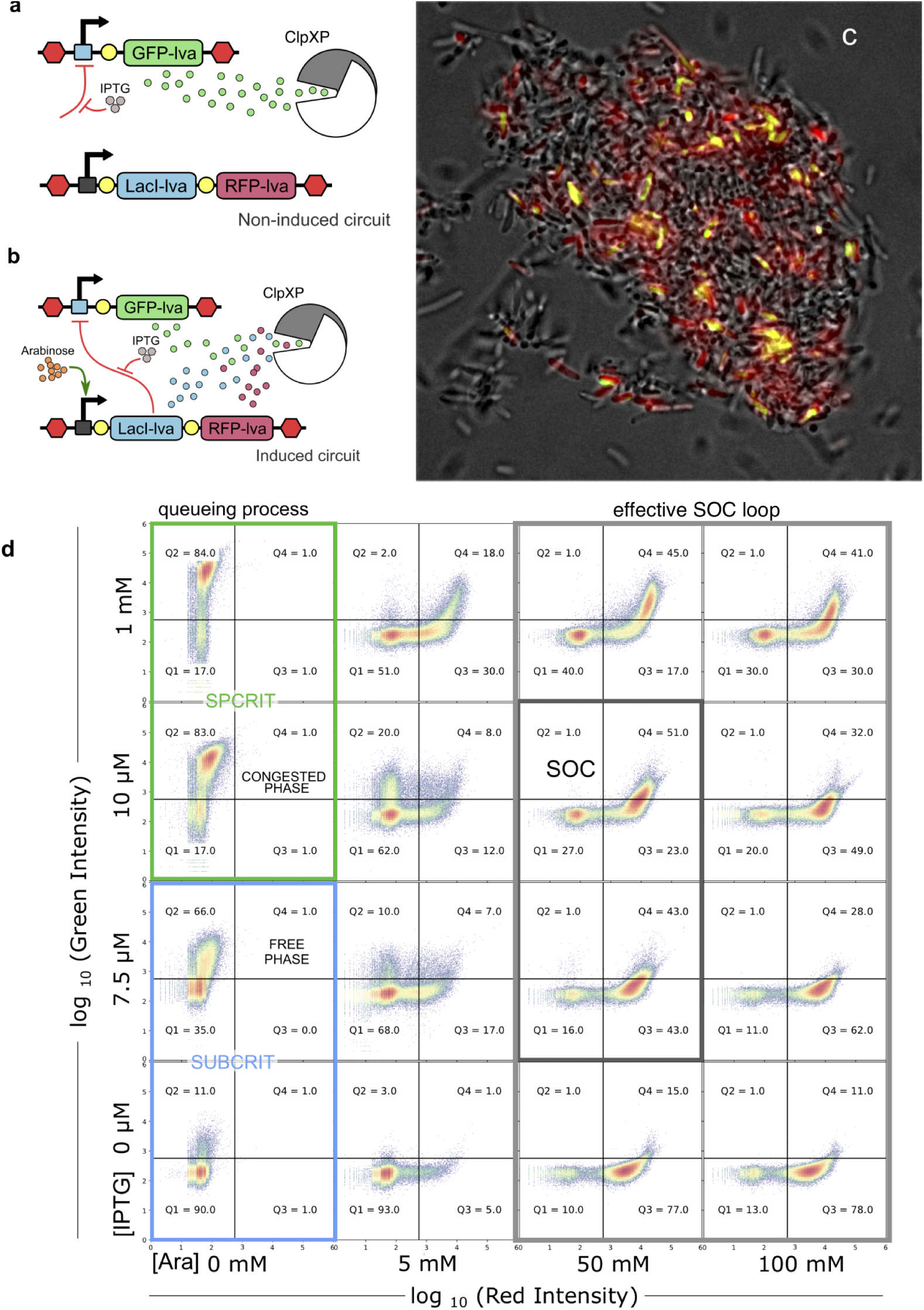
Engineered gene circuit implementing the SOC motif in *E. coli* cells. (a) The gene construct design in the non-induced state. The expression of ClpXP degradable GFP under the placI promoter can be regulated by the IPTG concentration present on the medium. (b) The gene construct SOC design in its induced state. The Arabinose (*Ara*) concentration controls the production of degradable LacI repressor and RFP, both, under the pBAD promoter. This design allows to use the red fluorescence as a proxy of LacI concentration. (c) Overlapped bright field and fluorescence images of bacterial culture induced with *Ara* (100 mM) and IPTG (10*µ*M). Here, yellow bacteria expresses both GFP and RFP. (d) Flow cytometry dot plots (green channel vs red channel) of *E. coli* cultures exposed to different concentrations of IPTG and *Ara*. In the non-induced circuit (a), without *Ara*, the transition from non-congested proteolytic machinery phase (i e free, SUBCRIT) to congested phase (SPCRIT) depends on the tunable GFP-lva production. As IPTG increases, hence the de-repression of GFP expression, ClpXP is not able to degrade the excess of GFP and cells are mostly with high emission in green. When the circuit is induced (b), *Ara* triggers the expression of the LacI repressor and the RFP reporter, that are also degraded by the ClpXP complex. The increase of tagged proteins to be degraded contributes to the congestion of ClpXP but also the LacI repression helps to de-congest by reducing the tagged GFP expression. Thus, as *Ara* concentration is increased, we notice a shift towards higher RFP levels along with a dispersal and lower levels of GFP values. This defines the domain (gray window) where the feedback loop required for SOC can effectively operate. A SOC state is obtained in the presence of high *Ara* concentration around the IPTG values close to the queueing transition. Most cells (Q3 + Q4 *≈* 80%) are emitting in the red channel, but exhibit broad range of green fluorescence levels, since this state is characterized by fluctuations associated to large bursts of GFP expression (the heterogeneous GFP expression is apparent in the yellow cells of (c) and in the histogram of Fig. 4). Flow cytometry analysis of more IPTG-*Ara* combinations are shown in Section III.D, SM.

ClpXP is responsible for degrading proteins carrying the SsrA or YbaQ degron sequences, reducing the half-life of a tagged protein from hours to minutes. In an exponentially growing *E. coli* culture (Optical Density (OD) OD_660_ from 0 to 2), the endogenous levels of ClpX and ClpP are constant and involve around 100 ClpXP molecules, which can degrade at least 10^5^ molecules of GFPssrA per cell per replication cycle. However, due to the limited number of ClpXP protease complexes, the degrading capacity of ssrA-tagged proteins can be easily saturated by over production of a synthetic tagged protein [49–51].

The non-induced circuit depicted in Fig. 3a is thus a particular instance of our more complex motif (Fig. 3b) that would correspond in our case to the presence of endogenous LacI, with the GFP-lva expression being repressed. The presence of Arabinose leads to a strong expression of the repressor LacIlva (our control parameter) and the reporter protein RFP-lva. Only when high levels of the -lva tagged proteins are reached, and the degradation machinery is saturated, there is enough LacI-lva to repress the production of GFP-lva. The repression loop is consistently removed when the production of GFP-lva (our order parameter) is reduced enough as for desaturate the ClpXP protease that can degrade the repressor again. The addition of Isopropyl *β*-d-1-thiogalactopyranoside (IPTG), as indicated by a negative input to the repression feedback (see Fig. 3b), switches on our SOC circuit and allows to control the level of GFP-lva expression. High levels of IPTG will lead to an overproduction of GFP-lva and the subsequent ClpXP complex congestion, thus reproducing the limit case that would correspond to the standard phase transition of the queueing process [26] (Fig. 1a-b).

The SOC motif is contained in a single, high-copy plasmid to ensure a maximal concentration of the vector while maintaining the parity of the two parts of the circuit (Fig. 3a-b). The construct was transformed in XL1-Blue *E. coli* strain. Further details of the cloning process and sequences can be found in Section S.III, SM. Also, this design allows an easy tuning to obtain parameters involving SOC, in terms of the strength of the promoter (i. e. pBAD) and by adjusting the efficiency of the repressor (in our case, IPTG for LacI, see Section III, SM).

To perform the experiments, a single colony was inoculated in a volume of 4 ml, and grown at 37°C until the exponential phase was reached with an OD_660_ around 0.6. This homogeneous fresh culture was then used to inoculate all the conditions used in the experimental design. Each combination was inoculated with 1 *µ*l of the starter culture to a final volume of 4 ml. Cells were grown for about 10 hours at 37°C, until reaching an approximate OD_660_ of 0.8-1. The output of the each condition was then analysed using both Fluorescence-Activated Cell Sorting (FACS) and fluorescence microscopy (Fig. 3c-d).

The results from the FACS are displayed in Fig. 3c, where a 4 × 4 array of different combinations of IPTG and Arabinose concentrations define our parameter space by means of dot plots. The range of concentrations shown here are 0 ≤ mM [IPTG] ≤1, 0 ≤ mM [*Ara*] ≤100 (see the specific values in Fig. 3c and in Section III, SM). As described above, these small molecules allow to explore a parameter space where we can move from a decoupling between the two genes to a full-fledged repression feedback required for criticality to occur. The different cell population responses to the tuning of both IPTG and *Ara* reveal the relative impact of each on the SOC motif. The target for a SOC state implies two requirements: (i) the expression of large enough levels of the control parameter to effectively perform its feedback; and (ii) a GFP expression characterised by bursts but displaying a low average activity. In the non-induced state, without arabinose (left column), increasing levels of IPTG concentration promote a standard transition from the free to the congested phase (sub- and super-critical phases, indicated in the two bottom and at the two top panels as SUBCRIT and SPCRIT, respectively). As IPTG grows, we effectively weaken the strength of the repression loop until a critical point is reached allowing congestion to rise. This is clearly observed from the displacement of the density dot plots from low to high levels of GFP.

As Arabinose concentration increases, we move in the other dimension of our parameter space, where the control molecule gets more common (cells emit in the red channel) but cannot always effectively act as a repressor. This clearly is a time point picture shot of a bacterial population that exhibits the fluctuating GFP levels characteristics of the SOC state. The same SOC behaviour is shown in the fluorescence microscope image of Fig 3c, where some bacteria do not have the ClpXP saturated (and thus do not display fluorescence), many are near the critical state of ClpXP saturation with lower levels of effective LacI-lva to repress the GFP, and exhibiting a wide range of GFP-lva concentrations (bacteria in yellow) and few bacteria have enough Laci-lva to degrade the GFP (bright only in red).

The experiments with *E. coli* confirm SOC fluctuations of GFP-lva, while the reporter of the control element (i. e. RFP-lva) remains basically stable in terms of the concentration levels and their dispersal. The experimental system successfully reproduces another important feature of criticality, namely the presence of a power law in the dynamics of the order parameter *σ*_1_. From the existence of a transition in the queueing process between the two phases described above, we can conjecture that the SOC motif should easily organize our gene network into this critical boundary provided that the control of the dimer concentration allows the loop to work properly. This is precisely what we found, as shown in Fig. 4a, where we display the same set of Arabinose-IPTG combinations shown in Fig. 3d. Here the statistics of expression are shown in Fig. 4a using cumulative distributions. The histograms of the non-induced *E. coli* colony with low Arabinose but tuned using IPTG (left panels) reveals a single-peak shape (a flat part followed by rapid decay in the cumulative plot). In general, as we increase the levels of both arabinose and IPTG, the distribution of our order parameter becomes fat-tailed once RFP levels become high, and well-defined power laws can be observed, as the two highlighted in Fig. 4b-c (the insets are linear-log plots to highlight the different behaviour of the two components of the SOC motif). A detailed parameter exploration is provided in Figs. S10-S17, SM.

**FIG. 4:**
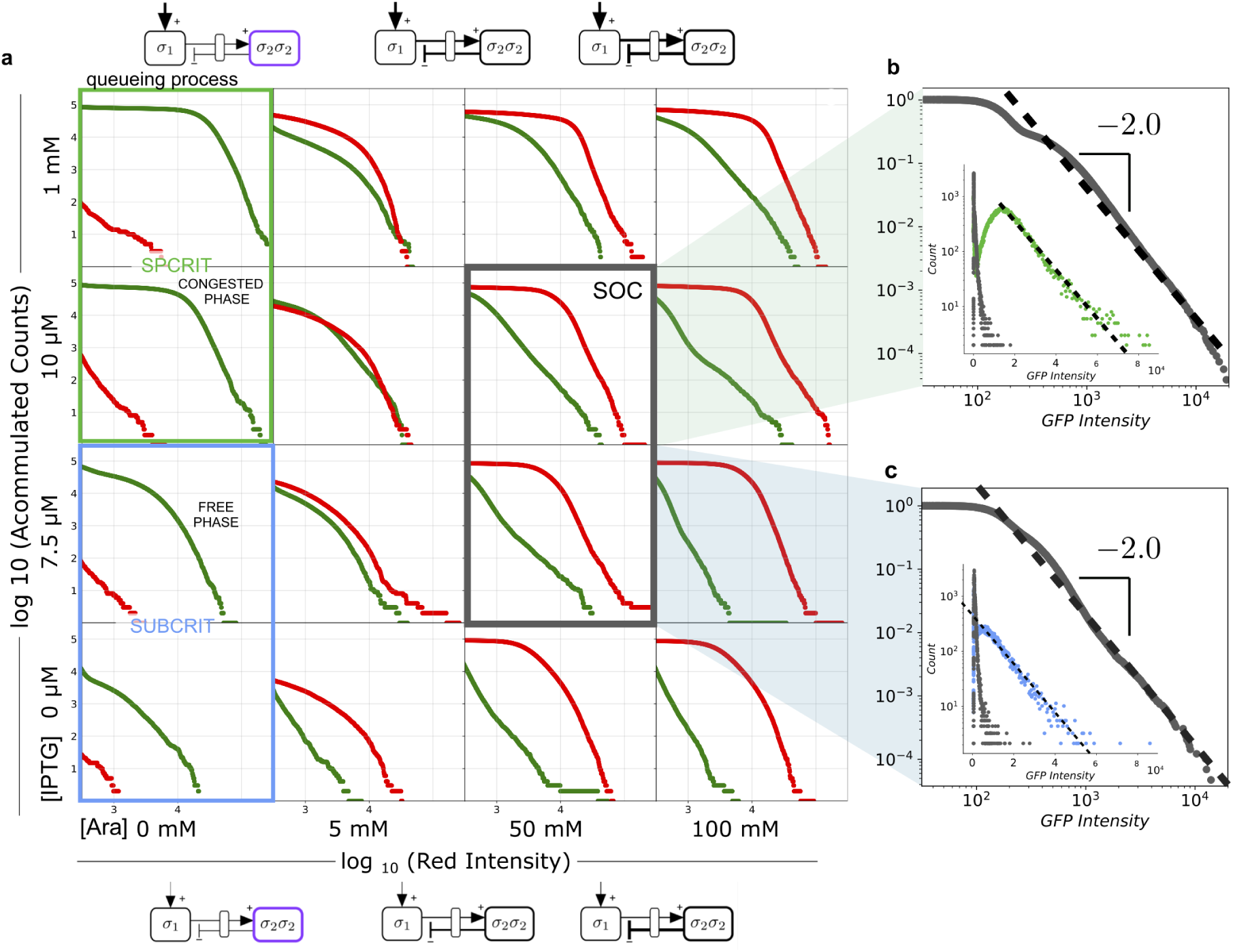
(a) Cumulative (non-normalized) distributions *P*_*>*_(*σ*_*i*_) of GFP and RFP fluorescence levels, here plotted using green and red lines, respectively, for the same set of conditions shown in Fig. 3d. The candidate combinations leading to the SOC state (grey panels) are characterized by a broad range of GFP expression revealed by the tail associated to large bursts. In (b) and (c) two cumulative histograms are shown for (10*µ*M IPTG, 50 mM arabinose (*Ara*)) and (7.5*µ*M IPTG, 50 mM *Ara*), respectively. Both distributions are close to a scaling law 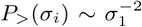 thus leading to a scaling exponent *γ* ∼3, consistent with the stochastic simulations. The insets display the comparison of the raw histograms of the congested state (green dots (b)) and free-phase (blue dots,(c)), respectively and the SOC state (gray dots) corresponding to the same IPTG conditions. Additional distributions with a more detailed IPTG-*Ara* combinations are shown in Section III.D, the SM.

## III. DISCUSSION

Self-organized criticality (SOC) has a seemingly paradoxical nature: it involves steady states that are always on the edge of instability. Are there intracellular processes poised close to critical points? Traffic dynamics in other contexts suggests that optimal flows occur close to criticality along with very broad fluctuations exhibiting optimal flow at critical points [22, 23]. Within cells, theoretical work suggests that enzymatic networks might be poised to criticality when the substrate input rate is balanced by the processing capacity of the enzymatic network [35] and that SOC states might pervade optimal growth [37]. Such critical balance would be a source of adaptation. In this paper we have followed a constructive approach by building a new type of network motif implementing the logic of SOC processes on a two-gene network by following the basic design principle of linking order and control parameters [31]. As the activity level (*σ*_1_) grows due to an overloaded proteolytic machinery, the competition for the ClpXP pool also increases the levels of the control component *σ*_2_ that can dimerize to perform a negative control on “emission” of *σ*_1_ thus effectively reducing activity.

Using the SOC motif architecture, it was possible to create a separation of time scales driving to highly fluctuating, critical dynamics. This work shows for the first time that this class of “unstable attractor” can be engineered in living cells. Interestingly, this is similar to the behaviour of computer traffic models where packet production is regulated by the amount of actual congestion [38], which leads to fat-tailed distributions of packets. Being at the critical state has important consequences linked with optimality and might be relevant for information-processing tasks. Several authors early suggested that biological computation could occur close to phase transitions [19, 39] and given the potential effects of a critical motif on other cellular systems performing given tasks, our results could give support to this conjecture at the cellular level. In this context, Hasty and co-workers have shown that a proper engineering of the proteolytic machinery can be used to achieve relevant functionalities, including tunable post-translational coupling [40] or *in vivo* drug delivery based on pulses of bacterial lysis against colorectal tumors [41]. Our SOC motif could further enhance some of these applications (wide fluctuations and rapid responses to external signals). An obvious extension of the critical motif could be a multicellular circuit able to trigger population-level avalanches by exploiting the quorum sensing machinery. Similarly, the fat-tailed behaviour could be wired to a diverse range of functionalities, such as search paths with fat-tailed statistics where bursting dynamics have adaptive value [43, 44].

Critical states are known to be part of the cognitive equipment of multicellular organisms, from the simple, non-neural placozoans to neural systems and animal collectives [45, 46]. The SOC motif might be a very efficient way of generating the highest phenotypic diversity in a microbial population and can be relevant to expand space of synthetic biology computational designs [47] into collective intelligence [48]. A missing point here is the lack of a time dimension that could help confirming our results and further develop a theoretical framework. This can be achieved by constructing a similar SOC motif within an eukaryotic cell, where the time scale of the resulting time series would be smaller than the cell division cycle. Finally, given the analogies between our system and critical traffic in parallel computer networks, an extension of our approach could involve a 3D spatially explicit system and the development of statistical physics models of critical intracellular traffic.

## IV. METHODS

### A. Plasmids construction

Plasmid construction and DNA manipulations were performed following standard cloning techniques. The LacI-lva (BBa_C0012), RFP-lva (BBa_K1399001) and GFP-lva (BBa_K082003) genes were amplified from the parts registry collection (2016). The forward primers were synthesized to contain the proper promoter and/or RBS sequences: the pBAD promoter and the RBS30 for the LacI gene, RBS34 for RFP gene and pLacIQ promoter with RBS34 for GFP gene. The PCR products pBad-RBS30-Lacil-va and RBS3-RFP-lva were joined together by assembly PCR, and cloned to pBluescript plasmid in the restriction sites EcoRI and XbaI. The PCR product pLacIQ-RBS34-GFP-lva was cloned to a Bluescript plasmid by SpeI and PstI. The resulting plasmids were joined together by ScaI and the blund ends of Eco53kI and EcoRV. The clonings were realized in the pBluescript II SK(+) plasmid backbone (ColE1 high copy number replication origin). See sequence of primers in the supplementary table I (see also Fig. S9, SM).

### B. Strains and growth conditions

Plasmid cloning and evaluation of the circuit behaviour was performed in *E. coli* XL1-Blue strain. All characterisation experiments were done in lysogeny broth (LB) Lennox media (10 g/L Tryptone 5 g/L Yeast Extract, 5 g/L NaCl) with a final ampicillin concentration of 125 *µ*g/mL. Single colonies were inoculated in 4 ml and grown at 37°C with shaking (200 r.p.m.) during 4 hours, to reach an approximate OD_660_ of 0.6. One microliter of the culture was re-inoculated in 4ml of fresh media, supplemented with ampicillin, and the corresponding Arabinose and IPTG concentrations. The cultures were grown overnight (10-14 hours) at 37°C with shaking.Once they were at OD_660_ of 0.8-1, were used for fluorescence measures.

### C. Imaging of single cell gene expression

The output of the SOC circuit was analyzed after 10h of incubation at 37°C with different combination of inputs. Samples were diluted in PBS and analyzed using flow cytometry (BD LSRFortessa). A total of 104 cells were collected from each sample. Specific emission fluorescence channels for GFP (FITC-H) and RFP (PE-H) were measured. A proper gate to subtract the debris particles was set using forward and side scattering channels. For the FACS graphics, the GFP and RFP fluorescence of cells inside the gate were plotted in adjacent axes. The cumulative distributions depict all bacteria with a FITC-H expression above 10^2.5^. All data was analysed and plotted using FlowJo (v7) software and customized Phyton code. The regression line and slopes of the histograms were calculated using Numpy and ploted with Matplotlib. For microscopic images, the cells were harvested at the same time than the cytometry analysis and pictures were collected with a inverted microscope Leica DMI6000, using a 40x oil objective. Bright field, red and green fluorescent images were taken, and then merged using ImageJ.

### D. Mathematical modelling

The mathematical model used here is a two-dimensional system of nonlinear ordinary differential equations describing the coupling between the order (*σ*_1_) and the control (*σ*_2_) parameters required to obtain criticality:

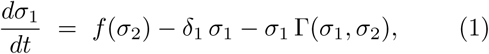

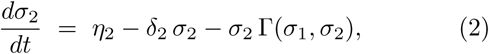

where the following Hill function response [33] is used:

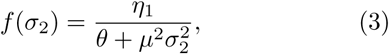

for the repression mediated by *σ*_2_*σ*_2_ dimers. The parameter *µ* ∈ [0, 1] weights the effect of IPTG on the strength of the negative control. When *σ*_2_ is small (the ClpXP system is working far from congestion) we have a production rate *f* (*σ*_2_ → 0) ≈ *η*_1_*/θ*. The inhibition function has a threshold value *θ* representing the concentration 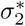 at which the rate drops to half its maximum value i. e. 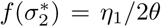. For larger values, it rapidly decays to zero. The saturation function, namely

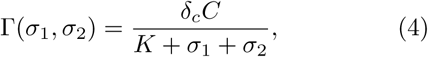

introduces the competition of both proteins for the proteolytic machinery. Here, as well, the limit case when no congestion occurs (due to low concentrations of both *σ*_1_ and *σ*_2_) gives a constant removal rate proportional to the concentration of ClpXP units, i. e. Γ(0, 0) = *δ*_*c*_*C/K*. The expression of *σ*_1_ gives the behaviour of the GFP-lva, whereas *σ*_2_ stands for LacIlva. Thus *f* (*σ*_2_) is the expression for the response of *pLac* (the promoter controlling the expression of GFP) to the LacI protein. In this function, *η*_1_ is defined as the production rate, *θ* the promoter sensitivity, and finally, *µ* weights how effective is the repression of LacI (effectiveness being altered by IPTG: the more IPTG the lower the *µ* value). The production rate of LacI (*σ*_2_) is controlled by the pBaD promoter, which will trigger a heavier production when there is Arabinose in the medium: the more Arabinose, the higher the value of *η*_2_. Both proteins are diluted and degraded at rates *δ*_1_ and *δ*_2_ respectively. Finally, both proteins are degraded by the ClpXP system, that can be saturated if there are enough proteins to be degraded. For this reason there is a sigmoid function, with degradation rate *δ*_*c*_, *C* standing for ClpXP concentration and sensitivity *K*. Notice that both proteins compete for the degradation machinery, thus inhibiting each other (being added in the denominator). Stochastic simulations of the previous deterministic model where also implemented using the Gillespie method [32] (see Section II.B, SM).

## Acknowledgments

The authors thank Jordi Garcia-Ojalvo as well as the members of the Complex Systems Lab for fruitful discussions. Special thanks to Arianna Bruguera for her help in the experimental implementation. RS thanks S. Kauffman, S. Manrubia and the late Per Bak for many discussions on criticality. This work was supported by the Botín Foundation by Banco Santander through its Santander Universities Global Division, the Spanish Ministry of Economy and Competitiveness, grant PID2019-111680GB-I00, a MICIN grant PID2019-111680GB-I00, an AGAUR FI 2018 grant, and the Santa Fe Institute (where the key idea was conceptualized). JS has been partially funded by the CERCA Programme of the “Generalitat de Catalunya”, by “Agencia Estatal de Investigación” grant RTI2018-098322-B-I00 and by the “Ramón y Cajal” contract RYC-2017-22243. AG has been funded by the AGAUR grant 2017-SGR-1049 and by the MINECO-FEDER-UE grants PGC-2018-098676-B-100 and RTI2018-093860-B-C21.

